# Dynamics of TFIIH and Spt4/5 during the transition from transcription initiation to elongation

**DOI:** 10.1101/2025.11.17.686592

**Authors:** James R. Portman, Kai Torrens, Gabriela G. Giordano, Jongcheol Jeon, Johnson Chung, Larry J. Friedman, Jeff Gelles, Stephen Buratowski

**Affiliations:** Department of Biological Chemistry and Molecular Pharmacology, Harvard Medical School, Boston, MA 02115; Department of Biochemistry, Brandeis University, Waltham, MA 02454

**Keywords:** RNA polymerase II, TFIIH, Spt5, DSIF, single-molecule microscopy

## Abstract

The transition of RNA polymerase II from initiation to elongation requires the timely exchange of many factors. Some initiation factors and elongation factors contact the same RNA polymerase II surfaces in an apparently mutually exclusive manner. For example, space occupied by the initiation factor TFIIH and elongation factor Spt4/5 (also known as DSIF) are predicted to overlap. To determine whether Spt4/5 actively displaces TFIIH, or binds only after TFIIH dissociates, labeled yeast nuclear extracts were combined with single-molecule fluorescence microscopy to observe their dynamics in real time. TFIIH categorically dissociates before Spt4/5 arrives, with a clear interval in between with neither factor present. Therefore, Spt4/5 binding is not required to dissociate TFIIH from RNA polymerase II.

**Significance Statement:** Proper regulation of eukaryotic gene expression is critical for proper development and response to the environment. Regulation often occurs during the transcription initiation and elongation stages of RNA synthesis. Here we use single-molecule microscopy to image the initiation factor TFIIH and the elongation factor Spt4/5, revealing their relative dynamics and the time-scales of their interactions with RNA polymerase II. We see that TFIIH dissociates before Spt4/5 arrival, indicating that the elongation factor is not required for initiation factor release from the polymerase.

## Introduction

Eukaryotic gene expression can be controlled at multiple steps, but regulation of transcription initiation and early elongation is common at most genes. Understanding the dynamics of transcription factors during these early steps is therefore essential. Both proteomics (1) and structural studies (2) indicate that major factor exchanges occur as RNA polymerase II (RNApII) pre-initiation complexes (PICs) transition to elongation complexes (ECs). For example, overlap in the positions of the initiation factor TFIIF and the Paf1 complex elongation factor (EF) suggest their interactions with RNApII may be mutually exclusive. Competition between TFIIF and the elongation factor ELOF1 regulates metazoan pausing (3). RNApII surfaces occupied by TFIIE, TFIIH, and Mediator in PICs also interact with Spt4/5 (DSIF in metazoans) in ECs (**Fig. 1A**) (4-7). Even in an archaeal transcription system, the TFIIE and Spt5 homologs compete for binding to RNA polymerase (8).

**Figure 1.**
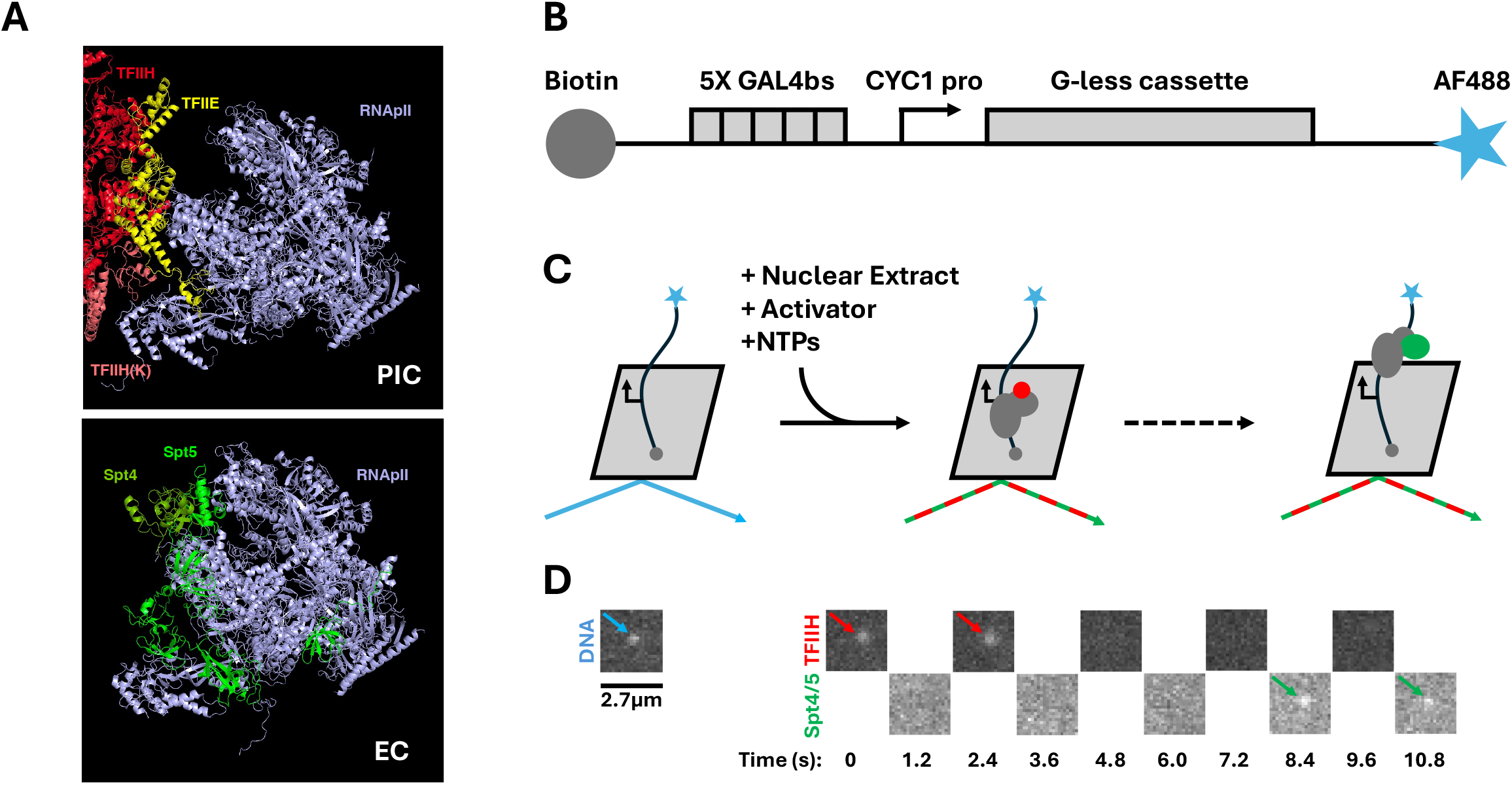
The CoSMoS system for imaging PIC to EC transitions. (**A**) Structures of the RNApII (blue) pre-initiation (upper panel: PIC: Protein Data Base ID 7LBM (6)) and elongation (EC: PDB ID 6TED (7)) complexes, showing the clash between binding of Spt4/5 (dark and light green) and the initiation factors TFIIE (yellow) and TFIIH (core module in red and kinase module in salmon). Note that nucleic acids and other factors in the structures are omitted. (**B**) Schematic diagram (not to scale) of the DNA transcription template bearing five Gal4 binding sites (bs), the *CYC1* core promoter (pro) and G-less cassette, with a biotin moiety at one end and an AF488 fluorophore at the other. (**C**) Outline of experimental procedure. DNA molecule locations are imaged using the blue laser; then nuclear extract, activator and NTPs are added; Kin28-HALO^JF646^ (marking TFIIH in the PIC) and Spt5-SNAP^DY549^ (marking Spt4/5 in the EC) are imaged by alternating excitation using the red and green lasers, respectively. (**D**) Example images (1s frame duration and 0.2s switching time) from a single DNA location; arrows denote detection of a fluorescence spot.

Two models for initiation to elongation complex transition are compatible with observed structures. The simplest is that the initiation factors must dissociate before the elongation factors can interact with RNApII. Early NTP-dependent events such as DNA unwinding and RNApII phosphorylation can trigger release of TFIIH and Mediator, and the growing RNA transcript in Initial Transcribing Complexes (ITCs) clashes with TFIIB in the RNA exit channel. These NTP-powered movements may liberate the RNApII surfaces needed for EF binding. In the alternative model, one or more EFs could initially bind RNApII while initiation factors are still present. Many EFs are large and modular, with multiple contact points on RNApII (4, 5). The incoming EFs might first contact non-overlapping surfaces on RNApII, and the partially bound EFs subsequently displace initiation factors by competing for the overlapping surfaces. A recent paper (2) argued for this displacement model based on *in vitro* experiments.

Here we use multi-wavelength colocalization single-molecule spectroscopy (CoSMoS) (9, 10) to observe and measure the dynamics of RNApII transcription factors in real time (11, 12). Our system is based on *S. cerevisiae* nuclear extracts, which contain the full complement of nuclear proteins needed for initiation and elongation (1). Using genetically encoded tags (13), two proteins of interest are fluorescently labeled with different colors, and labeled extracts are added to a microscope slide bearing tethered DNA template molecules. By imaging each protein at a rate of roughly one frame per second, the resulting data reveals association and dissociation rates, relative arrival orders, and kinetically distinguishable sub-populations. To test the two models proposed above, we imaged the PIC to EC transition.

Our CoSMoS experiments previously revealed that multiple basal transcription factors, along with RNApII and Mediator, have two modes of interaction with promoter DNA. In addition to the promoter-bound PIC, these factors assemble into a “Pre-PIC” that is tethered to activators at the Upstream Activating Sequence (UAS) (11, 14). Murakami and colleagues reported a cryo-EM structure likely to correspond to this intermediate (15). We have proposed that the Pre-PIC transfers to the core promoter for incorporation into the PIC. Importantly, TFIIH is not in the Pre-PIC, instead associating only at the core promoter as the final factor needed to complete the PIC (11). Furthermore, TFIIH crosslinks at promoters but not downstream transcribed regions (16, 17), and it is not detected in proteomic studies of isolated ECs (1, 18-20). Therefore, the presence of TFIIH on the DNA acts as a definitive marker of PIC presence.

RNApII ECs were detected by observing Spt4/5, which associates with RNApII during elongation (12). The Spt4/5 complex stimulates transcription elongation in multiple ways. The Spt5 NusG-like N-terminal domain binds across the RNApII cleft to stabilise more processive RNApII conformations, thereby reducing RNApII pausing and inhibiting template release, while its KOW and CTR domains can interact with and stabilise association of other elongation factors (4, 18, 21). Consistent with competition between initiation and elongation factors for RNApII binding, chromatin immunoprecipitation for Spt4/5 and other elongation factors show occupancy rising within the first few 100 bp of the gene body and remaining high throughout, anti-correlating with initiation factors (16, 17, 22, 23).

CoSMoS experiments in this study unambiguously show that TFIIH dissociates before Spt4/5 association with template DNA. Like the other basal factors (11), TFIIH is stable within PICs for up to several minutes in the absence of NTPs, while NTPs significantly accelerate dissociation, presumably after PICs begin transcribing. Spt4/5 binding to DNA templates was only seen in the presence of NTPs, as expected for an elongation factor. Simultaneous occupancy of both TFIIH and Spt4/5 on the same RNApII was essentially never seen. Instead, we observed a mean time interval of ∼35 seconds between TFIIH dissociation and Spt4/5 association. Our results therefore support a model where PIC disassembly occurs before elongation factors can bind RNApII.

## Results

### A CoSMoS system for imaging the PIC to EC transition

To study the conversion of PICs into ECs using CoSMoS, we constructed a *S. cerevisiae* strain in which the Kin28 (Cdk7 in metazoans) subunit of TFIIH had a C-terminal HALO tag and the Spt5 subunit of Spt4/5 carried a C-terminal SNAP tag. Nuclear extract was prepared in which Kin28-HALO was labeled with the JF646 fluorophore (24) and Spt5-SNAP was labeled with the DY549 fluorophore (New England Biolabs S9112S). These two dyes are excitable using the red and green laser, respectively. The transcription activity of this dual-labeled nuclear extract was confirmed in a bulk *in vitro* assay (**Sup. Fig. 1A**). Fluorescence imaging of a protein gel (**Sup. Fig. 1B**) showed a single labeled species corresponding to Spt5-SNAP^DY549^. Kin28-HALO^JF646^ also showed a major species at the expected size, although ∼25% of the HALO tag was observed as a released cleavage product, reducing Kin28 labeling efficiency.

The ∼650 bp transcription template carries a biotin at one end for attachment to the streptavidin-coated glass slide surface, and an AF488 fluorophore at the other end for imaging using the blue laser. The template contains five Gal4 binding sites (UAS), the *CYC1* core promoter, and a 300 bp G-less cassette (**Fig. 1B**). Each microscope field of view contained several hundred DNAs, and their locations were mapped before extract addition, together with a small number of fluorescent beads that act as fiducial markers to correct for stage drift during the experiment. Labeled nuclear extract, NTPs, and Gal4-VP16 activator were added to the flow chamber, and TFIIH and Spt4/5 were imaged using alternating excitation with the red and green lasers. The start of imaging is defined as time 0, and the cycling time through both channels was roughly 2.4 seconds (**Fig. 1C**). The arrival, dwell, and departure of individual TFIIH and Spt4/5 complexes were monitored for 45 minutes, during which hundreds of binding events were typically observed.

Images from the red and green channels were analyzed for fluorescent spot colocalization with DNA locations, using a set of proximity and intensity thresholds. A control set of off-target locations were used to determine the level of non-specific binding to the slide surface (as well as to locations where the dye on the DNA was photobleached or absent). TFIIH and Spt4/5 fluorescent spots were largely colocalized with DNA sites, and their “time to first binding” curves for the DNA sites fit well to a single-exponential DNA-specific binding model (**Fig. 2A, B** and **Table 1**). The initiation-to-elongation transition should be NTP-dependent, so the same CoSMoS experiment was performed without NTPs. TFIIH still bound to DNA locations far more frequently than off-target locations (**Fig. 2C**). The calculated association rate (*k*_specific_) was comparable to that with NTPs present (**Table 1**), consistent with a lack of NTP requirement for TFIIH incorporation into the PIC. In contrast, Spt4/5 showed essentially no binding to DNA locations in the absence of NTPs (**Fig. 2D, Table 1**), as expected in the absence of ECs. Similar results were obtained when frequencies were calculated using all binding events (**Table 2**). Unexpectedly, both on- and off-target Spt4/5 binding were far lower in the absence of NTPs, possibly hinting at an NTP-dependent modification or activation.

**Table 1.** Association rates for on-(*k*_specific_) and off-target (*k*_non-specific_) locations, with and without NTPs. Ranges represent standard errors calculated from 1,000 bootstrap samples. Values of *k*_specific_ and k_non-specific_ for Spt4/5 absent NTPs could not be determined (n.d.) because there were too few binding events (see **Table 2**). *A*_f_ represents the active fraction of DNA molecules; n represents number of DNA or off-target locations monitored.

|  | Time to First Binding Fits |  |  |
| --- | --- | --- | --- |
|  |  | +NTPs | -NTPs |
| TFIIH | $k_{\text{specific}} \text{ (s}^{-1}\text{)}$ | $4.9 \pm 0.9 \times 10^{-4}$<br>(n=202) | $8.0 \pm 2.2 \times 10^{-4}$<br>(n=322) |
| | $k_{\text{non-specific}} \text{ (s}^{-1}\text{)}$ | $0.32 \pm 0.04 \times 10^{-4}$<br>(n=408) | $0.42 \pm 0.06 \times 10^{-4}$<br>(n=368) |
| | $A_f$ | $0.56 \pm 0.06$ | $0.28 \pm 0.05$ |
| Spt4/5 | $k_{\text{specific}} \text{ (s}^{-1}\text{)}$ | $5.5 \pm 2.9 \times 10^{-4}$<br>(n=202) | n.d.<br>(n=322) |
| | $k_{\text{non-specific}} \text{ (s}^{-1}\text{)}$ | $0.92 \pm 0.09 \times 10^{-4}$<br>(n=408) | n.d.<br>(n=368) |
| | $A_f$ | $0.35 \pm 0.11$ | $0.01 \pm 0.01$ |

**Table 2.**
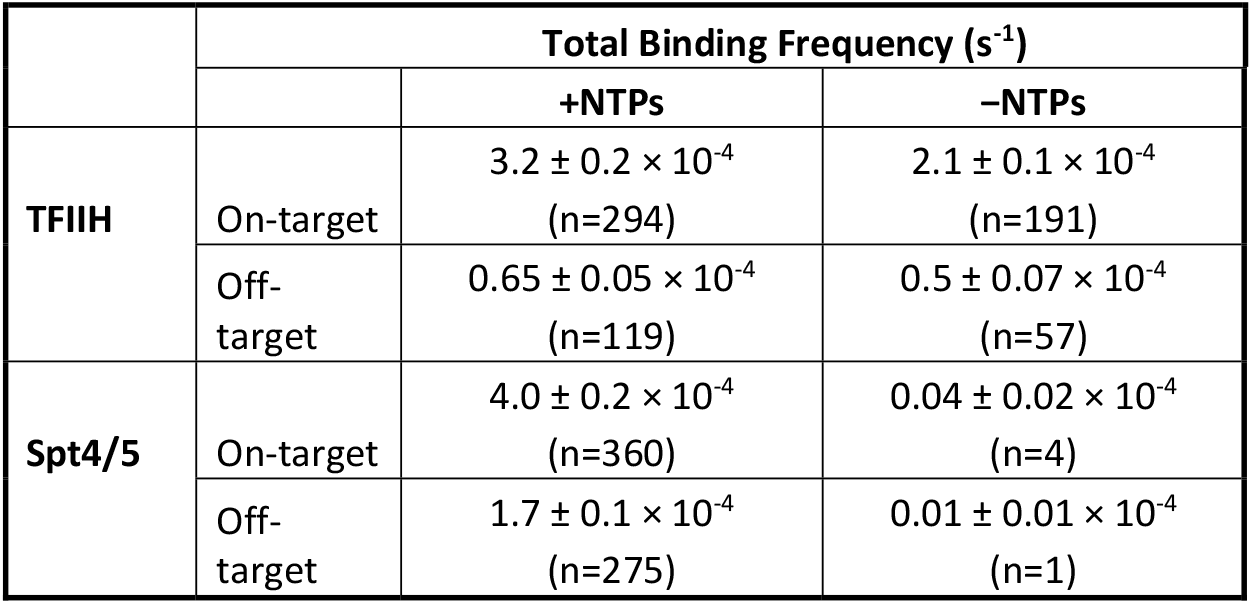
Total binding frequency for on- and off-target locations with and without NTPs. Errors represent SE. n represents number of colocalization events.

| | Total Binding Frequency ($\text{s}^{-1}$) | | |
| --- | --- | --- | --- |
|  |  | +NTPs | -NTPs |
| TFIIH | On-target | $3.2 \pm 0.2 \times 10^{-4}$<br>(n=294) | $2.1 \pm 0.1 \times 10^{-4}$<br>(n=191) |
| | Off-target | $0.65 \pm 0.05 \times 10^{-4}$<br>(n=119) | $0.5 \pm 0.07 \times 10^{-4}$<br>(n=57) |
| Spt4/5 | On-target | $4.0 \pm 0.2 \times 10^{-4}$<br>(n=360) | $0.04 \pm 0.02 \times 10^{-4}$<br>(n=4) |
| | Off-target | $1.7 \pm 0.1 \times 10^{-4}$<br>(n=275) | $0.01 \pm 0.01 \times 10^{-4}$<br>(n=1) |

**Figure 2.**
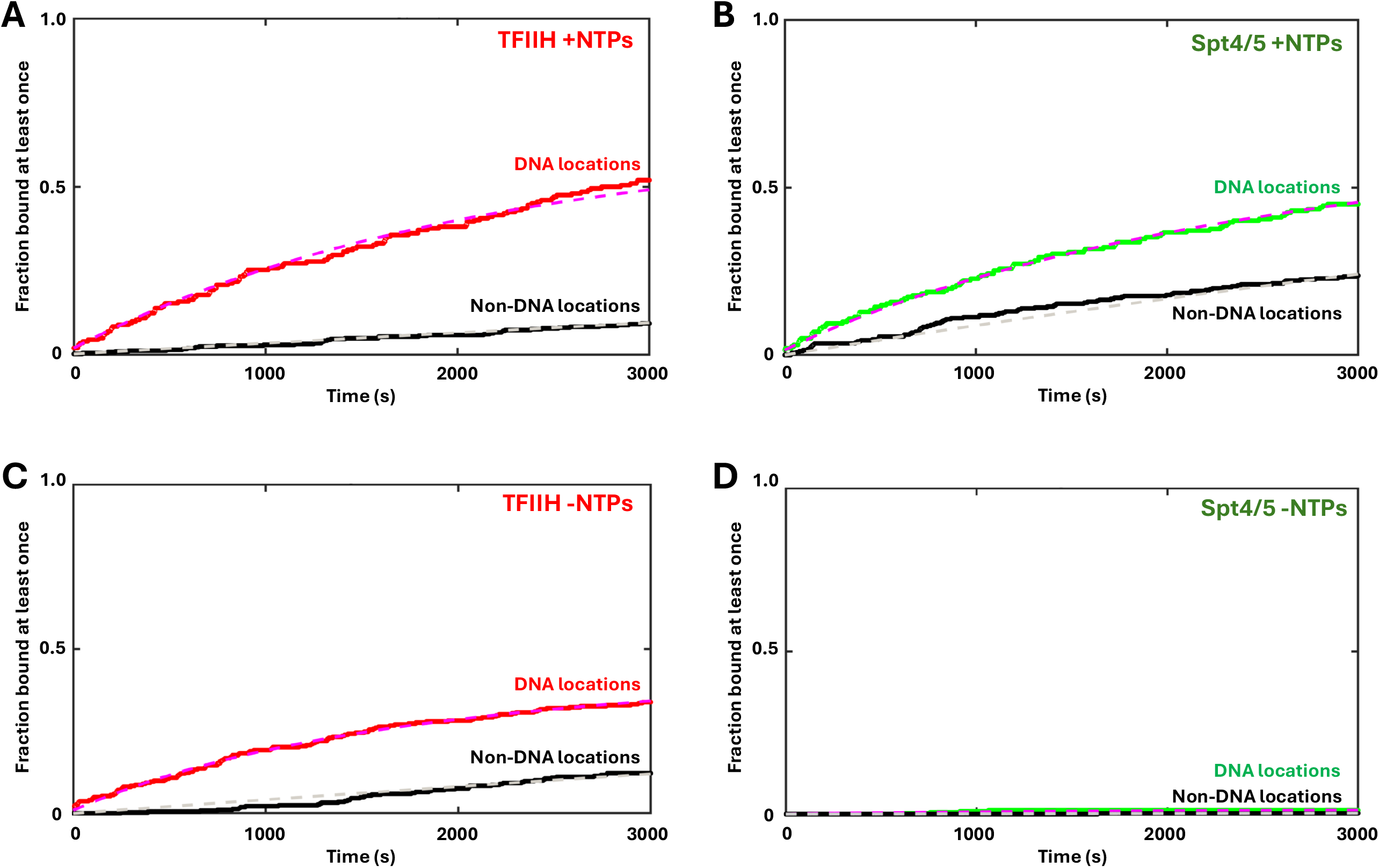
Measurement of TFIIH and Spt4/5 association with template using CoSMoS. “Time to first binding” curves represent the cumulative fraction of on-target (color) or off-target (black) locations showing colocalization of the fluorescent protein (red for TFIIH and green for Spt4/5). Reactions were performed either in the presence (**A** and **B**) or absence (**C** and **D**) of NTPs. Dashed lines show curves from fitting data to a single exponential DNA-specific binding model (see **Table 1** and Methods).

### TFIIH dissociates before Spt4/5 arrival

Intensity traces show factor dynamics at single DNA locations, reflecting the behaviors of the initiation and elongation complexes (see four representative examples in **Fig. 3A**). Only one TFIIH complex at a time was observed on each DNA, representing a single PIC assembling at the core promoter. Spt4/5 was also typically seen at one molecule per template, consistent with elongation complex structures showing a single Spt4/5 complex. Occasionally, an additional intensity jump representing a second Spt4/5 was seen, which likely occurs when a new EC is formed before the earlier one has dissociated from the template. To visualize the collective reaction, intensity traces from 100 randomly chosen “active” DNA locations (i.e., those bound by TFIIH at least once during the experiment) were converted to a binary function and plotted as a rastergram (**Fig. 3B**). Data from each active DNA is represented using a single horizontal ribbon, with horizontal colored bars spanning intervals during which colocalized protein spots are detected. The ribbons are stacked by order of first TFIIH arrival.

**Figure 3.**
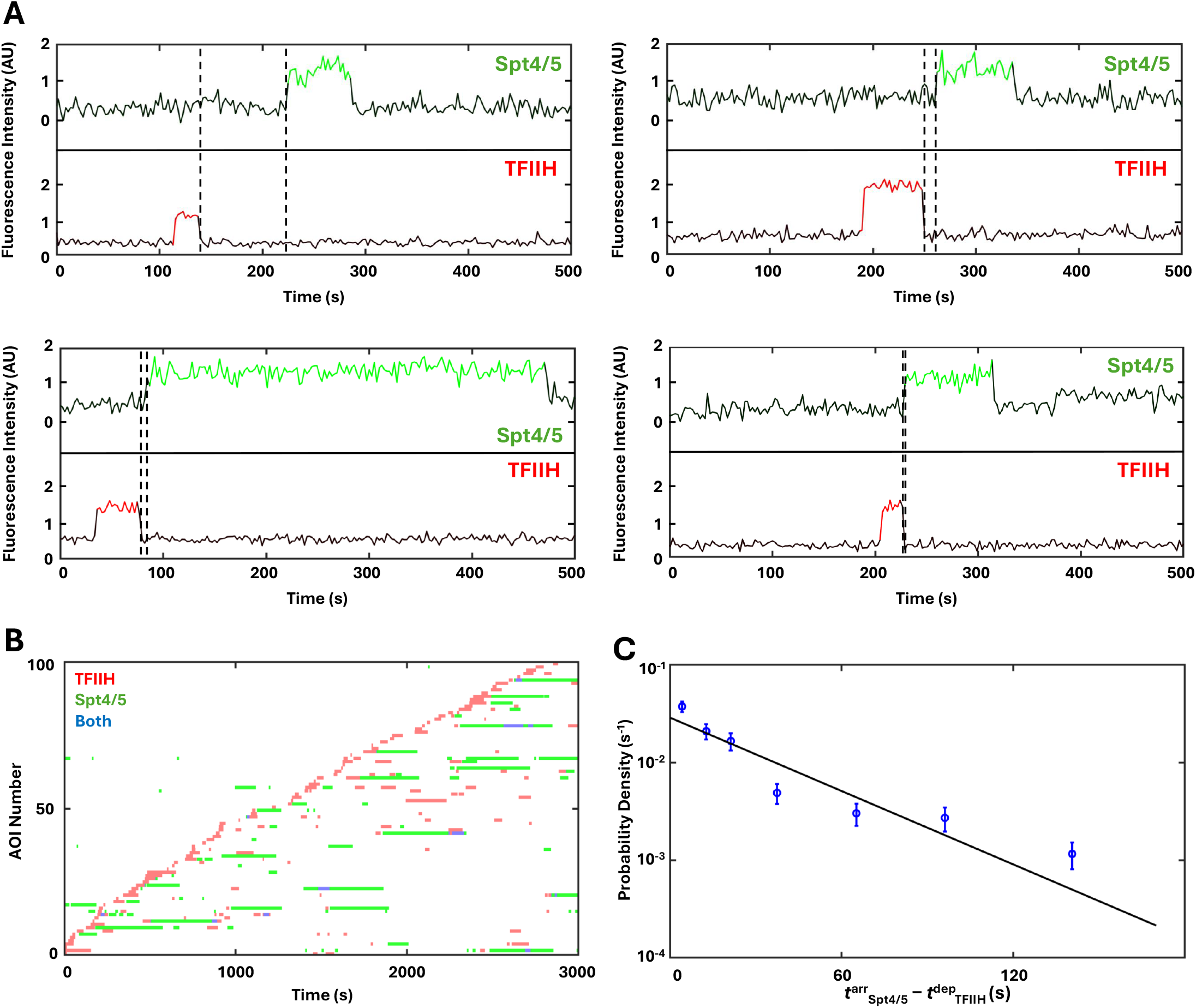
Single-molecule imaging of TFIIH and Spt4/5 binding. (**A**) Fluorescence intensity time records at four representative DNA locations, coded according to the presence (color) or absence (black) of a colocalized TFIIH (red) or Spt4/5 (green) fluorescence spot. Dashed lines mark times between TFIIH departure and Spt4/5 arrival. (**B**) Dual-channel rastergram of 100 randomly chosen DNA locations (AOI, Area of Interest) that showed TFIIH binding, sorted by time of first TFIIH arrival. Each horizontal row represents a single DNA location, and the bars represent protein residence intervals. Red marks the presence of TFIIH, green the presence of Spt4/5, and blue the presence of both proteins. See **Table 2** for association frequencies calculated from all binding events. **(C**) Distribution (±SE) of the time intervals between TFIIH departure (t^dep^_TFIIH_) and Spt4/5 arrival (t^arr^_Spt4/5_). Line shows maximum-likelihood fit to a single-exponential function (n=146).

Both individual traces (**Fig. 3A**) and the rastergram (**Fig. 3B**) show that Spt4/5 binding was rarely seen unless TFIIH had already visited the same DNA. The first protein to arrive – 77 ± 2.7% of the time (181 of 235 DNAs) – was TFIIH. Spt4/5 binding events before TFIIH likely represent either non-specific slide surface binding near the DNA, or transcription events primed by TFIIH that was unlabeled, photobleached, or cleaved from the HALO tag (see **SI Appendix Fig. S1B**). Unsurprisingly, not all PICs progressed through elongation, with 41 ± 3.7% (74 of 181 DNAs) of TFIIH binding events never being followed by Spt4/5. These events may reflect abortive initiation or premature termination. However, Spt4/5 binding after ∼59% of TFIIH binding events suggests a relatively high efficiency of PIC to EC conversion in our nuclear extract system relative to other estimates from in vivo (25) and in vitro studies (26-28), which typically report efficiencies in the range of 1-10%.

By looking at DNA molecules bound first by TFIIH, we could determine whether Spt4/5 arrives whilst TFIIH is still present, consistent with a mechanism in which Spt4/5 facilitates basal factor dissociation; or instead Spt4/5 arrives after TFIIH has departed – indicating that TFIIH, and potentially other basal transcription factors, must dissociate before Spt4/5 can bind. The results in **Figure 3** categorically show that Spt4/5 binds only after TFIIH has already departed. Individual traces always show a gap where neither protein is present (**Fig. 1D** and **Fig. 3A, B**).

There were a very few observations of apparent colocalizations between TFIIH and Spt4/5 (blue sections in **Fig. 3B**). These fell into two classes, each with less than five occurrences. In the first, TFIIH appeared when Spt4/5 was already bound. These events likely occur when a new PIC forms at the promoter while an EC is still on the DNA. Traces of this type show a TFIIH binding event before the Spt4/5 arrival and subsequent overlap between Spt4/5 and a second TFIIH arrival. A less likely explanation is overlapping non-specific binding of Spt4/5 and TFIIH at the same off-target location. In the second class, the overlap lasted only a single frame. Inspection of the intensity traces and raw images shows that these are rastergram artifacts that occur when TFIIH dissociation and Spt4/5 binding occur in rapid succession during a single red-green imaging cycle (**SI Appendix, Fig. S2**).

To quantitatively probe the dynamics of the initiation-to-elongation transition, data from three replicate CoSMoS experiments were pooled. We identified 146 events where Spt4/5 arrived within 170s after TFIIH departure. Intervals longer than that were excluded as being unlikely to be biologically relevant. These events were not simply due to coincidence, as random pairing of the Spt4/5 and TFIIH traces from different DNA molecules produced only 25 instances of Spt4/5 binding within 170s of TFIIH. The distribution of time intervals between TFIIH departure and Spt4/5 arrival (*t*^arr^_Spt4/5_ − *t*^dep^_TFIIH_) was fit by a single exponential function with a mean of 35 ± 4 (SE) s (**Fig. 3C**), presumably reflecting the time it takes Spt4/5 to find the newly available binding surface on the EC. This time interval is less than the 71s we previously observed between template associations of Pol II and Spt4/5 (12), consistent with the interpretation that RNA polymerase II recruitment to the template is followed by TFIIH binding and release, which is in turn followed by Spt4/5 recruitment.

TFIIH consists of two subcomplexes: the core module contacts RNApII and DNA to mediate ATP-dependent promoter melting, while the kinase module phosphorylates RNApII to regulate its interactions. Because these modules can function separately in certain contexts (29-32), it was important to test if departure of Kin28 coincides with dissociation of the entire TFIIH complex. Accordingly, a nuclear extract having both kinase (Kin28-SNAP^DY549^) and core (Rad3-HALO^JF646^) modules labeled was imaged. Consistent with the existence of free kinase module (29), there were frequent binding events of Kin28-SNAP^DY549^ without Rad3-HALO^JF646^. A smaller number of Rad3-only events were also seen. However, the large majority of these single protein events lasted two frames or less, and similar binding was observed at sites without DNA. Using a threshold for binding lasting more than two frames (∼2.8 seconds), more than half of events showed co-binding of both proteins. For the vast majority of these events, the two proteins arrived apparently simultaneously, i.e. within one green-red imaging cycle (**SI Appendix, Fig. S3A**). However, the time distribution clearly showed events where one or the other protein arrived within 10 seconds of the other. This behavior suggests that some free core and kinase module may independently incorporate into the PIC to co-stabilize their binding, a possibility that will be pursued in the future. Relevant to this study, in the presence of NTPs, 82% of Kin28 and Rad3 co-binding events lasting longer than two frames showed simultaneous departure of both proteins (**SI Appendix, Fig. S3B**). Since simultaneous photobleaching is very unlikely, this high percentage of simultaneous departure indicates that any photobleaching occurs on a slower timescale than dissociation. In the minority of cases where one module does leave before the other, the remaining module departs within 8s – well below the mean time of 35s it takes Spt4/5 to arrive. We therefore conclude that Kin28 departure can be used as a marker of TFIIH dissociation.

### Mechanistic features revealed by TFIIH and Spt4/5 dwell times

Important information can be gleaned from the distribution of the factor dwell times, which are defined, for TFIIH, as the time from TFIIH initial colocalization with a DNA site (incorporation into the PIC) until TFIIH dissociation. The dwell intervals of TFIIH (n=624) in the presence of NTPs were plotted as a Probability Density Function (PDF). Plotting this data using a log scale vertical axis (**Fig. 4A**) is useful for revealing subpopulations, as the distribution shape can reveal the existence of different bound species, even if they represent a fraction of the total binding events (33). An exponential decay produces a straight line, so inflections from linearity, as seen at ∼200s, suggests the presence of at least two components. Accordingly, the TFIIH dwell data were fit to a bi-exponential function. The fast-dissociating component, representing 85% of events, had a mean dwell of 29 ± 6s. The slow component, representing 15% of events, had a mean of 5,000 ± 2,000s. The long TFIIH dwells that contributed to the slow component were rarely followed by Spt4/5 binding and likely represent PICs or TFIIH stuck in off-pathway states. Excluding these long dwells, the distribution made up only of TFIIH dwell times that led to Spt4/5 association (**Fig. 4A**, red points, n=135) are not kinetically distinct from the total population (**Fig. 4A**, blue points, n=623). This congruence is consistent with the notion that commitment to EC formation does not occur until after TFIIH departure.

**Figure 4.**
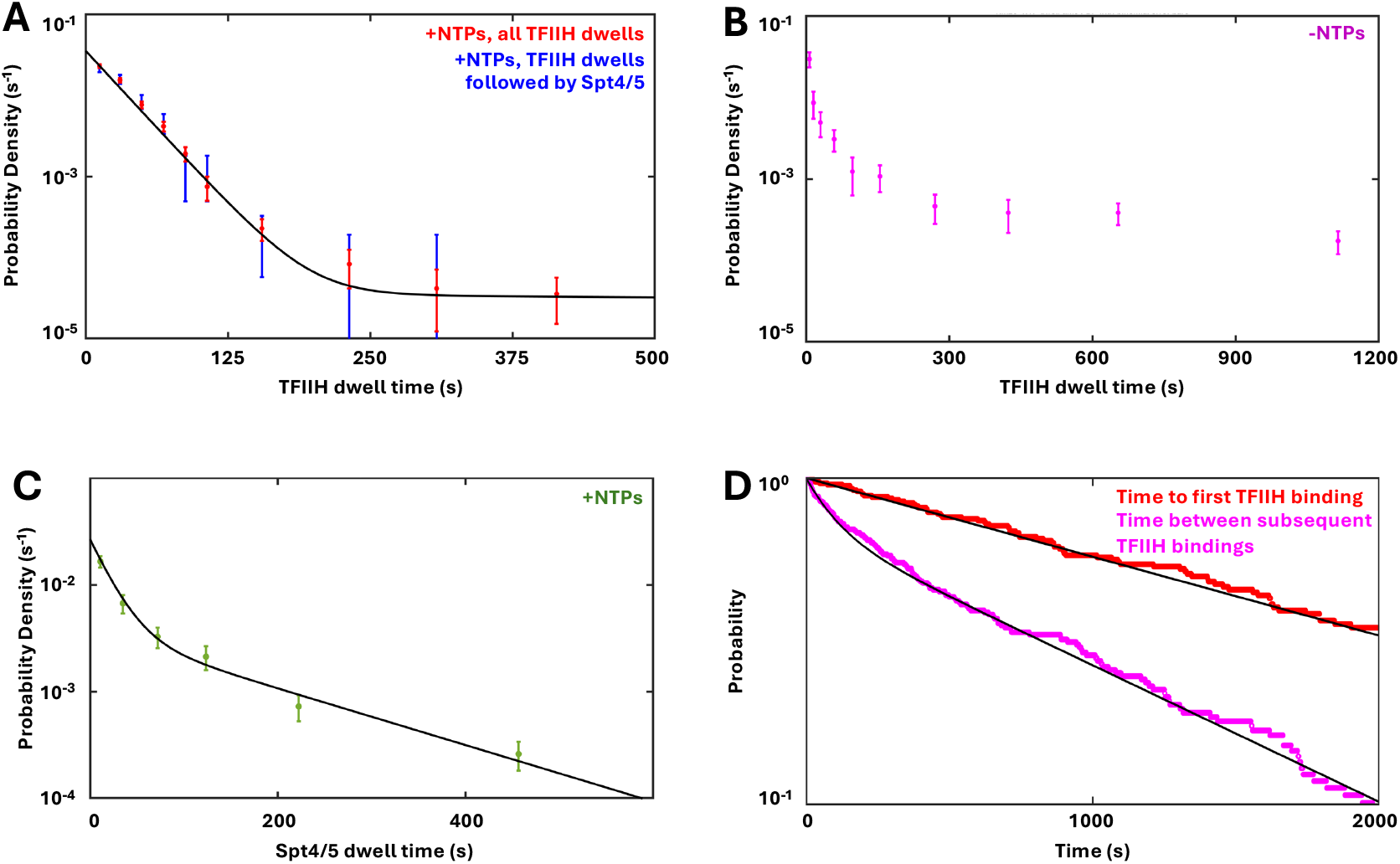
Dwell time distributions of TFIIH and Spt4/5. **A**) Probability density (±SE) of all TFIIH dwell times in the presence of NTPs (red; n=623) and of the subset of TFIIH dwell times that were followed by Spt4/5 binding within 170s (blue; n=135). Black line shows bi-exponential fit of all dwell times **B**) Distribution of TFIIH dwell times in the absence of NTPs (magenta; n=92). **C**) Distribution of Spt4/5 dwell times that follow TFIIH within 170s, with line showing fit to a bi-exponential function (n=117). **D)** Cumulative survival plots of times to initial TFIIH binding to each DNA (red; n=112) and times between subsequent TFIIH bindings (magenta; n=178). Black lines show the associated single- and bi-exponential fits, respectively.

In contrast to the reaction containing NTPs, the dwell time distribution of TFIIH in the absence of NTPs was not fit well by a biexponential and may contain more than two components (**Fig. 4B**). The fast component TFIIH dwells have a comparable lifetime but represent a smaller fraction than those seen in the presence of NTPs, which may be due to minimal residual ATP or NTPs in the nuclear extracts. However, there are many more longer TFIIH dwells, many of which likely represent fully-formed PICs awaiting NTPs to proceed to elongation phase.

The dwell time distribution of Spt4/5 is fit by a bi-exponential function (**Fig. 4C**). The fast component (accounting for 45% of events) has a mean duration of 21 ± 6s, similar to the 47 s lifetime reported previously (12). The slow component (accounting for 55% of events) has a mean duration of 160 ± 34s (n=117). This population may have escaped detection in the earlier work (12) because the DHFR tag used to label Spt5 in that study binds dye non-covalently, with the resulting dye dissociation limiting the duration of observable binding. The fast component of the Spt4/5 lifetime likely represent ECs that transcribe through the G-less cassette and run off the end of the DNA, as well as any ECs where Spt4/5 dissociates but RNApII remains. An EC transcribing the ∼300 bp template in 21 seconds produces an average rate of ∼14 nt·s^-1^, which compares well with speeds of ∼13 nt·s^-1^ recorded *in vivo* (34) and comparable to 17-40 nt·s^-1^ in single molecule studies using purified components (28, 35). The slow component of the Spt4/5 lifetime may therefore reflect a sub-population of ECs that remain bound to the DNA end (36).

### Evidence for accelerated reinitiation

While examining the rastergram (**Fig. 3B**) to analyze PIC-to-EC transitions, we noticed that TFIIH events on each DNA often appeared to be clustered together. This behavior can be quantitatively analyzed by comparing the distributions of times before the initial TFIIH binding with the times between subsequent TFIIH bindings (i.e. between the first TFIIH departure and second TFIIH arrival, departure of the second and arrival of the third, and so on). If each binding event is independent (i.e., requires repetition of the upstream rate-limiting steps), the shapes of the distributions should be similar. The distribution of times to first TFIIH binding fit well to a single-exponential function with a mean of 1,800 ± 400s, consistent with a single rate-limiting step (**Fig. 4D**; red data). In contrast, the distribution of times between subsequent TFIIH binding was clearly bi-exponential (**Fig. 4D**; magenta data). The long-lived component (representing 65% of the data) had a mean time of 1010 ± 100s, comparable to the mean of first TFIIH binding. However, for the remaining 35% there was only 105 ± 50s on average before a TFIIH molecule returned to the DNA. This result suggests some non-initial TFIIH binding events may take advantage of one or more factors remaining on the template after the previous TFIIH dissociates.

This faster-than-expected assembly of PICs is significant, as several mechanisms have been proposed to facilitate transcription reinitiation. *In vivo*, temporally clustered transcription is observable as “bursting” (37). A recent study imaging transcription *in vivo* showed that bursts correlated with periods of extended activator presence (38). *In vitro* experiments suggest that a subset of coactivators and/or basal transcription factors may remain associated with promoter DNA after the first initiation event, leading to faster re-assembly of subsequent PICs (27, 39, 40). Our finding that TFIIH dissociates before binding of Spt4/5 to the EC argues against a model where TFIIH is part of a reinitiation scaffold (27). Using CoSMoS, we observed that multiple pre-PICs can assemble on multiple activators bound at the UAS, leading us to propose that these produce bursting by sequential transfer from the UAS to the core promoter during windows of TBP occupancy (11).

## Discussion

Deciphering how essential transcription factors coordinate their timely association and dissociation from RNApII is important for understanding gene expression. Using the kinase subunit of TFIIH to mark PICs and Spt5 subunit of Spt4/5 to mark ECs, we imaged the transition between initiation and elongation complexes. Our single-molecule system, which combines nuclear extracts with CoSMoS, revealed several key kinetic details that were not possible to derive from ensemble *in vitro* assays or *in vivo* chromatin immunoprecipitation experiments.

First, TFIIH dwell times in the presence of NTPs suggest 29 ± 6s after TFIIH incorporation are needed for PICs to transition into elongation in our system. This time is consistent with another *in vitro* single-molecule system using optical tweezers and purified yeast factors, which observed RNApII movement along the DNA approximately 17 seconds after NTPs were added to pre-formed PICs. This period would encompass the time needed for TFIIH to push the downstream DNA into the RNApII active site and for transcript initiation on the unwound template strand. TFIIH likely remains associated with the initial transcribing complex, although some contacts with RNApII are already released (41).

A second key observation is that TFIIH binding precedes, but clearly dissociates prior to, Spt4/5 arrival. The lack of Spt4/5 signal in the absence of NTPs confirms its validity as a marker of elongation. The gap between occupancy by the two factors rules out a mechanism in which association of Spt4/5 is necessary to either actively or indirectly facilitate TFIIH removal from RNApII. This conclusion contrasts with a recent study positing that binding of the human homolog of Spt4/5 (DSIF) and the Negative Elongation Factor (NELF) facilitated displacement of TFIIH and other basal transcription factors from initial transcribing complexes (2). While a species-specific difference cannot yet be ruled out, it should be noted that the experiment underlying the proposed displacement model did not directly analyze PIC to EC transition. Instead, purified RNApII was pre-assembled with a DNA template and 24 nt transcript. These minimal EC scaffolds were then used for equilibrium binding competitions, in the absence of NTPs, between the basal and elongation factors. Although this reconstituted system has proven invaluable for creating the ECs needed for structural studies, the lack of other key factors and requirement for equilibrium conditions makes the results less reliable for inferring kinetic mechanisms of the PIC to EC transition.

Finally, our nuclear extract system exhibits behavior consistent with facilitated reassembly of PICs, which can contribute to transcription bursting (37, 42). While bursting is likely to occur by multiple mechanisms at multiple time-scales, a single-molecule system should prove especially useful for testing some of the models proposed to explain this phenomenon.

The nuclear extract system accurately reproduces most features of RNApII transcription initiation and elongation (1, 19, 43), but the dynamics may be slowed compared to the in vivo reaction. The 29 second average TFIIH dwell time measured in our system is longer than estimates from in vivo single-particle tracking, which report average chromatin residence times of 4-10 seconds for TFIIH subunits within the nucleus (44, 45). Here, the TFIIH 29 second average dwell time plus an average gap of 36 seconds between TFIIH departure and Spt5 arrival sums to ∼65 seconds, aligning well with our previous study estimating an average time of ∼71 seconds between RNApII arrival at the promoter and Spt5 arrival (12). Single-particle tracking experiments (45) estimated average dwell times of ∼43 seconds for RNApII and 20-25 seconds for Spt4 and other elongation factors, suggesting a maximum delay of 18-23 seconds between RNApII arrival and incorporation of Spt4-Spt5 in the elongation complex. The slightly slower dynamics in the in vitro system are likely because transcription factor and NTP concentrations are lower than in vivo, as well as the lack of super-coiling on the linear DNA templates. Nonetheless, the numbers from both systems are reasonably close, and together they provide time ranges useful for modeling factor dynamics.

Our single-molecule studies have provided other new insights into gene regulation not observed in ensemble assays. Imaging the interactions between transcription activators and the coactivators SAGA and Mediator revealed a simple mechanism for synergy between multiple activators at enhancers (14, 46). Unexpected kinetic behaviors of RNApII and the general transcription factors led to the discovery that some of these factors pre-assemble into a pre-PIC while tethered to activators at the UAS/enhancer (11, 12). We expect that future single-molecule experiments will uncover additional important features of gene expression.

## Materials and Methods

### Yeast strain and nuclear extract preparation

*S. cerevisiae* strain YSB3356 (*MATa, ura3-1, leu2-3,112, trp1-1, his3-11,15, ade2-1, pep4Δ::HIS3, prbΔ::his3, prc1Δ::hisG, kin28(L83G, V21C)* (1)) was transformed sequentially with two DNA integration cassettes made by PCR using our previously described template plasmids (13). One cassette fused three copies of the influenza hemagglutinin epitope (3xHA) and a SNAP tag to the C-terminus of Spt5 and was marked with a NatMX selectable marker. The second cassette C-terminally fused an HA3-HALO tag to Kin28, marked by KanMX. In each case, successful integrants were selected by their drug resistance marker, and in-frame expression and labeling of the fusion protein were verified by immunoblotting and fluorescent dye labeling. The resulting strain, YSB3770 (*MATa, ura3-1, leu2-3, 112, trp1-1, his3-11,15, ade2-1, pep4Δ::HIS3, prbΔ::his3, prc1Δ::hisG, kin28(L83G, V21C)-HA3-HALO::KanMX, Spt5-HA3-SNAP::NatMX*), was used to make fluorescently labeled nuclear extracts as previously described (11, 19).

The same approach was used to create yeast strain YSB3764 (*MATa, ura3-1, leu2-3, 112, trp1-1, his3-11,15, ade2-1, pep4Δ::HIS3, prbΔ::his3, prc1Δ::hisG, RAD3-HA3-HALO::KanMX, KIN28-HA3-SNAP::NatMX*), used for the experiments imaging Kin28 and Rad3. Yeast nuclear extracts were prepared and labelled as previously described (11, 46). By comparing protein fluorescence and the HA tag signal, dye labeling efficiencies were estimated to be at least 80-85% for both SNAP and HALO.

### DNA template preparation

The template for bulk *in vitro* transcription assays was pUC18_G5CYC1_G-(SB649), which carries five Gal4 binding sites and the CYC1 core promoter upstream of a cassette directing synthesis of a G-less RNA (47). The DNA template used for single-molecule microscopy experiments (**Fig. 1B**) was made by PCR from pUC18_G5CYC1_G-(SB649), amplified using a biotinylated forward primer (5′-/5Biosg/TTGGGTAACGCCAGGGT-3′) and a AF488-labeled reverse primer (5′-/5Alex488N/AGCGGATAACAATTTCACACAG-3′) (IDT). The PCR product was twice purified using DNA SizeSelector-I SPRI magnetic beads (Aline Biosciences Z-6001).

### In vitro transcription assay

Nuclear extracts were tested for transcription activity *in vitro* in a bulk assay. Typically, a 30 μl reaction consisted of 15 μl of nuclear extract, 100 ng plasmid DNA template (pUC18_G5CYC1_G-; SB649), 200 nM Gal4-VP16, 0.33U/μl RNAsin, and an ATP regeneration system comprising 93 ng/μl creatine kinase and 10 mM phosphocreatine. The reaction contained 20 mM HEPES pH 7.6, 100 mM potassium acetate (KOAc), 1 mM EDTA, and 5 mM magnesium acetate (Mg(OAc)_2_). ATP, UTP, GTP, and CTP were added to 400 μM of each to initiate transcription. After 40 minutes at room temperature, 200 μl of stop solution (300 mM NaCl and 5 mM EDTA) and 1000 units of RNAse T1 were added and incubated at 37°C for 20 minutes. Next, 13 μl of 10% SDS and 2.5 μl of 20 mg/ml proteinase K were added and the reaction incubated at 37°C for 30 minutes. Following this, 10 μl of 10 M ammonium acetate (NH_4_OAc) and 2 µl of 10 μg/μl yeast tRNA were added, and reactions then extracted with phenol-chloroform and ethanol precipitated. RNA was resuspended in 10 μl primer annealing mix (5 mM Tris pH 8.3, 75 mM KCl, 1 mM EDTA) with 1 pmol of fluorescent oligonucleotide (/5Cy5/TGATAGATTTGGGAAATATAGAAGAAGGA) that hybridizes to the G-less cassette. This mix was heated to 90°C for 1 minute then left to cool to 48°C over the next 45 minutes to allow the primer to anneal to the RNA. Next, 20 μl synthesis mix (50 mM Tris pH 8.3, 75 mM KCl, 4.5 mM MgCl_2_, 15 mM DTT, 150 μM each of dATP, dCTP, dTTP, dGTP and 100 units MMLV-Reverse Transcriptase) was added and incubated at 42°C for 30 minutes to allow reverse transcriptase to generate single-stranded DNA complementary to the RNA. Reactions were stopped by adding one-tenth volume 3 M NaOAc and three volumes of cold ethanol to precipitate cDNA. The pellet was resuspended in formamide loading buffer, electrophoresed on a 6% acrylamide 8 M urea denaturing gel, and imaged with an Amersham Typhoon. A set of activator-dependent bands corresponding to multiple transcription start sites in the CYC1 promoter are observed (**SI Appendix, Fig. S1A**).

### Slide preparation

Glass microscope slides (24 × 60 mm; (VWR 48404-133) and 25 × 25 mm coverslips (VWR 48366-089) were rinsed by sequentially spraying with distilled water and then 90 proof ethanol (VWR 89094-618) from wash bottles. They were then placed in glass containers (VWR 74830-150) containing ethanol, which were then placed inside a water bath sonicator (VWR 89375-452) and sonicated for 10 minutes at the maximum setting. Slides and coverslips were then rinsed by repeatedly filling and emptying the glass containers with deionized (DI) water to remove the ethanol. Slides and coverslips were then submerged in 1 M potassium hydroxide (KOH) and sonicated again for 15 minutes. This cycle of DI water rinsing and KOH sonication was repeated once. Following this, the slides were transferred to a plastic holder (VWR 82003-433) containing acetone (VWR EM-AX0115-1) to remove residual water, emptying the container, adding fresh acetone, and sonicating for 1 minute. After a third acetone rinse, a 1.5% Vectabond-acetone (Vector Labs SP-1800-7) solution was added for 5 minutes. Once treated with Vectabond, the slides and coverslips were briefly rinsed with distilled deionized water and dried using filtered air. All surfaces are visibly hydrophobic at this point (water droplet contact angle of > 100°).

For passivation and DNA linkage of slides, mPEG-SG-2K (Laysan Bio) and biotin-PEG-SVA-5K (Laysan Bio) were dissolved in filtered 0.1 M sodium bicarbonate, mixed at a 200:1 ratio with 25% w/v PEG overall, and ∼30 µl added to the center of each slide. Coverslips were placed on top of the slides such that both surfaces were exposed to the PEG mixture. They were then left in a closed container, together with wetted paper towels to maintain humidity, at room temperature for 3 hours. All surfaces were then washed with 10 mM Tris acetate (pH 8.0), dried with filtered air, and stored at -80°C in a sealed container backfilled with nitrogen gas. Just before each experiment, lanes and wells were drawn on the slide with silicone vacuum grease, and the coverslip was placed on top to create a simple flow cell with lane volume ∼20μl (9). After placing the assembly on the microscope stage for fluorescence visualization, streptavidin-coated fluorescent beads (ThermoFisher T10711; diluted by ∼262,000x) were added until 2-4 beads were present per field of view. These provide fiducial markers in each image to correct for stage drift or other motion. For slide application, beads were suspended in TPAGBT buffer: 50 mM Tris acetate (pH 7.9), 100 mM KOAc, 8 mM Mg(OAc)_2_, 27 mM ammonium acetate, 1 mg/ml BSA (Gold Biotechnology A-420-100), 0.1% Tween-20 (VWR EM-9480), 2% glycerol. After washing out unbound beads by twice flowing through 25μl of TPAGBT buffer, TPAGBT solution containing 0.01 mg/ml streptavidin was added for one minute and washed out just before adding TPAGBT solution containing 10 pM biotinylated fluorescent DNA molecules. After monitoring the fluorescence in the microscope for the desired number of bound DNA molecules, binding was terminated by washing with TPAGBT buffer containing an oxygen scavenging system (0.9 U/ml protocatechuate dioxygenase (Sigma P8297) and 5mM protocatechuic acid (Sigma 03930590)) and triplet state quenchers (1 mM Trolox (Sigma 238813), 1 mM propyl gallate (Sigma 02370), and 2 mM 4-nitrobenzyl alcohol (Sigma N12821)).

### Microscopy

Microscopy was performed with a micromirror total internal reflection fluorescence microscope (9). DNA images were taken by excitation with the blue (488 nm) laser at 1 mW power (all powers measured incident to the micromirror) just before adding the transcription reaction mixture into the flow cell. The reaction conditions were essentially the same as in the bulk *in vitro* transcription assay, except for addition of the oxygen scavenging system and triplet state quenchers (see above), 1 mg/ml BSA, 20 ng/μl *E. coli* genomic DNA digested with HaeIII, and 0.1% Tween-20. For experiments that omitted NTPs, the ATP regeneration system was replaced with an ATP-depletion system (20 mM glucose and 2 units hexokinase (Sigma H4502-2.5KU)) to deplete residual NTPs that may be present in the nuclear extract. Images were collected in each channel (red and green) every 2.4 s (1s exposure time per frame, plus 0.2 s switching times) for roughly 1 hour. Both the green (532 nm) and red (633 nm) lasers were used at a power of 0.5 mW. Custom software “Glimpse” implemented with LabView operated the microscope, laser shutters, filter wheels, camera, and image acquisition (https://github.com/gelles-brandeis/Glimpse).

### Image quantification and single-molecule kinetic analysis

Fluorescence images were analysed using the MATLAB-based program imscroll as previously described (9, 48). Colocalization of proteins with DNA positions was determined using a set of proximity and intensity thresholds. Time intervals when the protein of interest is bound or not bound at DNA or control off-target locations without DNA were defined, and “time to first binding” plots and rastergrams were generated (48). To verify the computational image analysis, all instances in which Spt4/5 arrived within a short time from TFIIH departure were validated by visual inspection of fluorescent images to confirm TFIIH was not detected after Spt4/5 appeared.

To measure the rate constant for non-specific binding to non-DNA locations, the time to first binding data at these locations was fit to an exponential probability distribution function using maximum-likelihood, yielding *k*_non-specific_. Time to first binding at DNA locations data were then fit to a function that included both exponential non-specific binding with the measured *k*_non-specific_, and exponential specific binding to an active subset of DNA locations, yielding the active fraction (*A*_f_) and the specific binding rate constant, *k*_specific_ (48). These values are listed in **Table 1**. In addition, total binding frequency was calculated by dividing the total number of times a protein was observed to bind at the measured locations by the sum of all time intervals during which the protein was absent at the same locations (**Table 2**). Total binding frequency uses all binding events, while time to first binding (48) uses only the initial binding events on each DNA to minimize possible complications from dye blinking.

Dissociation rates and frequencies were visualized by plotting dwell time probability density distributions or cumulative survival plots (48). Distributions were fitted to single or double exponential functions as noted. For the TFIIH dwell time distributions, one-frame events were excluded from the fitting due to potential under-counting. Intervals between departures and arrivals were analyzed similarly.

## Acknowledgements

We thank the members of the Buratowski and Gelles labs for advice and sharing reagents. JRP is a Gordon and Betty Moore Foundation Postdoctoral Fellow. This work was supported by NIH grants R01GM056663 to SB and R01GM081648 to JG.

## Notes

### Competing Interest Statement

The authors have declared no competing interest.

### Summary of Updates

Discussion was expanded in response to reviewer's comments. Figure 1A was modified, changing colors of the proteins to match the rest of the paper.

