## Supplemental Figures 1-3 for "Dynamics of TFIIH and Spt4/5 during the transition from transcription initiation to elongation"

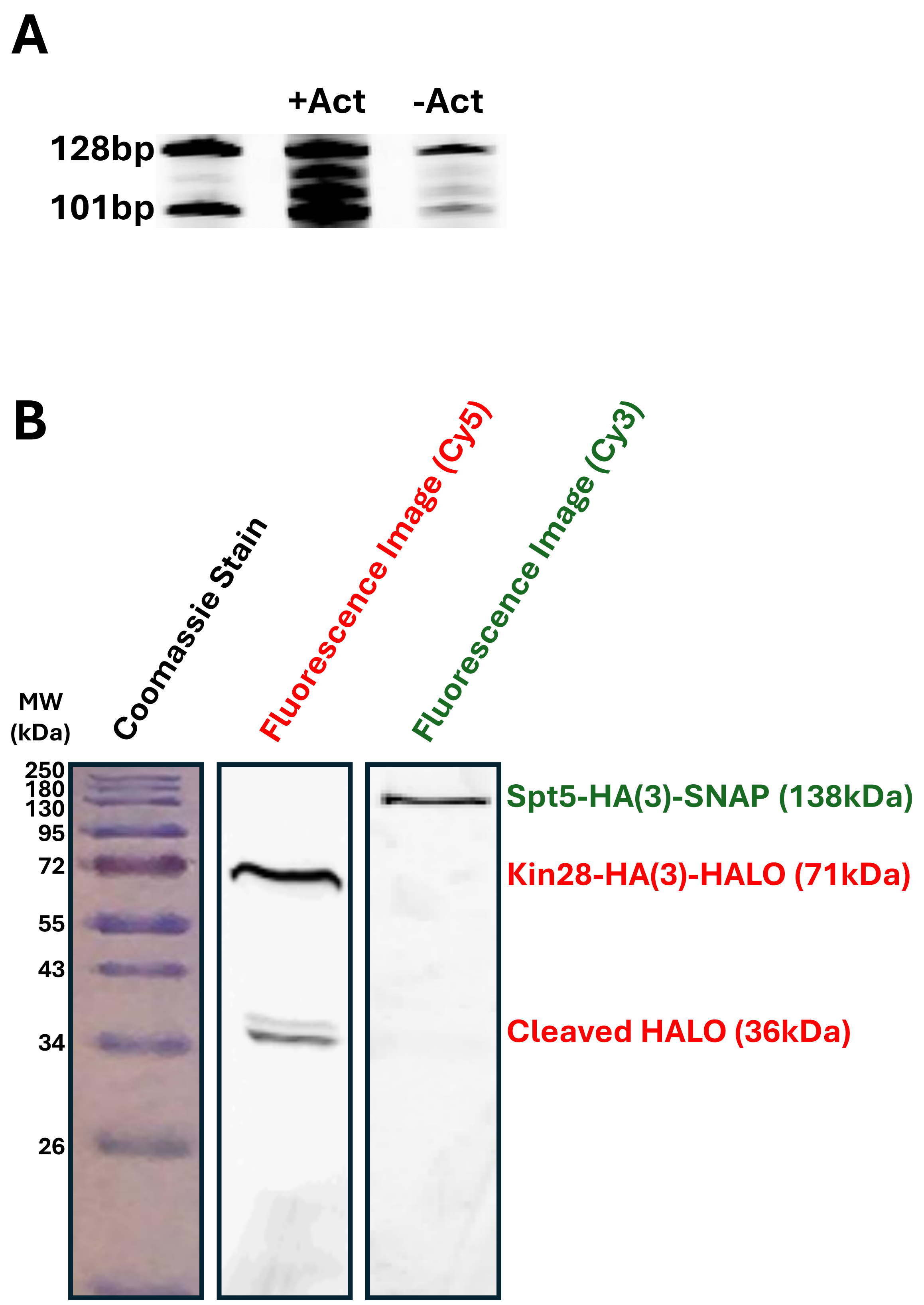
A
+Act
-Act
128bp
101bp
B
Coomassie Stain
Fluorescence Image (Cy5)
Fluorescence Image (Cy3)
MW
(kDa)
250
180
130
95
72
55
43
34
26
Spt5-HA(3)-SNAP (138kDa)
Kin28-HA(3)-HALO (71kDa)
Cleaved HALO (36kDa)

**Figure S1**. Validation of nuclear extract activity and labeling. **A**) Bulk *in vitro* transcription assay using fluorescently labeled YSB3770 nuclear extract. RNA produced in the reaction was analyzed by primer extension using a Cy3-labeled primer. Fluorescent cDNA was run on an 8M urea polyacrylamide gel and imaged in the Cy3 channel. The presence of Gal4-VP16 (+Act; Lane 2) strongly stimulates RNA transcription relative to the control (-Act; Lane 3). The cluster of bands from 101-128 bp (see Lane 1 markers) is due to multiple transcription start sites in the *CYC1* promoter. **B**) SDS-polyacrylamide gel analysis of labeled YSB3770 nuclear extract. Coomassie stained ladder (left) and fluorescence images in the Cy3 (green) and Cy5 (red) channels (right and center, respectively) of a 10% SDS-PAGE gel containing the same YSB3770 nuclear extract used in panel A and CoSMoS experiments. Fusion proteins are visible at the expected sizes with some cleavage product visible for Kin28-HA3-HALO.

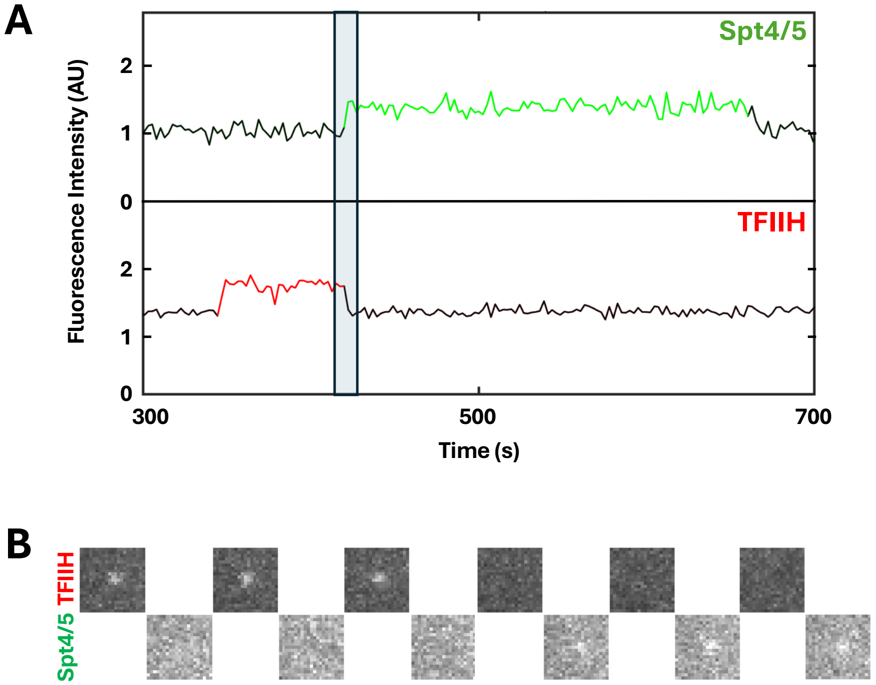

A
Spt4/5
TFIIH
Fluorescence Intensity (AU)
2
1
0
2
1
0
300
500
700
Time (s)
B
Spt4/5 TFIIH

**Figure S2.** Raw images from an event showing possible TFIIH-Spt4/5 co-occupancy. **A)** Intensity vs time trace of DNA AOI #22, which showed a possible overlap in the rastergram (**Fig. 3B**). Shaded box shows the region corresponding to individual frame images in panel B. **B.** Frame-by-frame images of the transition between TFIIH and Spt4/5 on DNA AOI #22 (as in **Fig. 1D**, with 1s frame duration and 0.2s switching time) during the transition. The microscopy images reveal that TFIIH departs just as Spt4/5 arrives.

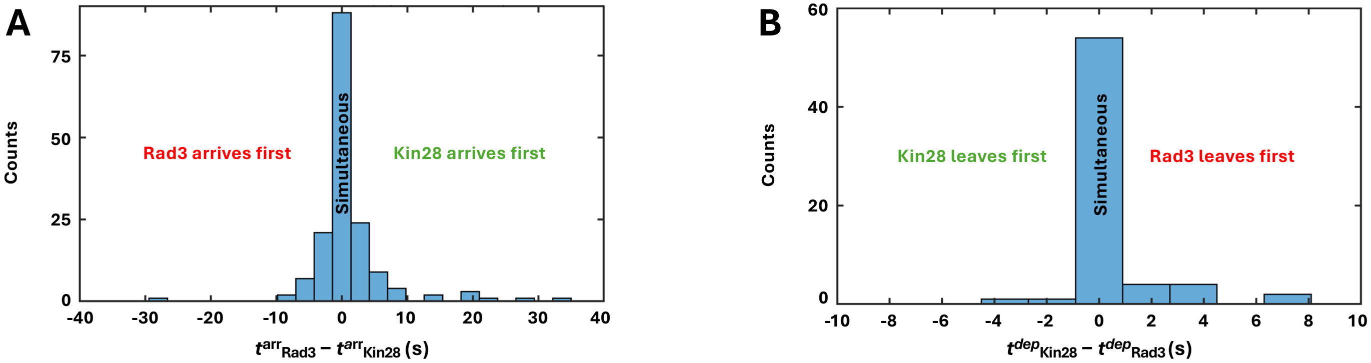

A
Counts
Rad3 arrives first
Simultaneous
Kin28 arrives first
0
25
50
75
-40
-30
-20
-10
0
10
20
30
40
$t^{arr}_{Rad3} - t^{arr}_{Kin28}$ (s)
B
Counts
Kin28 leaves first
Simultaneous
Rad3 leaves first
0
20
40
60
-10
-8
-6
-4
-2
0
2
4
6
8
10
$t^{dep}_{Kin28} - t^{dep}_{Rad3}$ (s)

**Figure S3.** Temporal coordination between TFIIH core and kinase modules. **A)** Histogram showing the distribution of times (n=164) between Rad3-HALO$^{JF646}$ and Kin28-SNAP$^{DY549}$ arrivals on DNA ($t^{arr}_{Rad3}$ – $t^{arr}_{Kin28}$). This data combines multiple experiments done both in the presence and absence of NTPs, which did not show differences in arrival behaviors. **B)** Histogram showing the distribution of times between Rad3-HALO$^{JF646}$ and Kin28-SNAP$^{DY549}$ departures from DNA ($t^{dep}_{Kin28}$ – $t^{dep}_{Rad3}$) in the presence of NTPs (n=66).
